# Dynamic geometry remapping of neural activity within frontal and subcortical areas during decision-making

**DOI:** 10.64898/2026.06.11.731612

**Authors:** Frederic M. Stoll, Neelima Valluru, Peter H. Rudebeck

## Abstract

Decision-making involves dynamic evaluation of competing options. Recording 16,495 neurons across nine frontal and subcortical areas in macaques revealed brain-wide fluctuations between encoding the attributes of available options. These dynamic representations track deliberation and scale with decision difficulty. Instead of binding attribute information of different options together, our analyses show that these representations are associated with dynamic area- and attribute-specific changes in neural geometry.

---

Decision-making is a dynamic process. When deciding between different options we often change our minds as we compare different attributes such as reward probability, sensory identity, and other potential costs. This deliberative process depends on distributed cortico-limbic circuits. Single neurons across prefrontal cortex (PFC) subdivisions, amygdala, and striatum encode attributes associated with different choice options during value-based decision-making (Padoa-Schioppa and Assad, 2006; Rudebeck et al., 2008; Kennerley et al., 2009; Stalnaker et al., 2014; Costa et al., 2019; Stoll and Rudebeck, 2024a, 2024b). Closer examination of activity within frontal cortex during deliberation has revealed a rich temporal structure: rather than encoding options simultaneously, populations of neurons dynamically switch between representations of each choice option (Rich and Wallis, 2016; Balewski et al., 2023; Panichello et al., 2024). These dynamic patterns of neural activity have primarily been characterized in individual parts of the frontal cortex, raising the question of whether other frontal and subcortical areas exhibit similar temporally-specific representations. Beyond characterizing their prevalence, a key open question is what computational role such dynamics play. One possibility is that these neural population activity patterns act as an attentional spotlight (Maunsell, 2015), boosting coding of whichever option is currently represented through a relatively uniform gain increase. Alternatively, these patterns could actively reshape how other decision-relevant attributes are encoded through population geometry reconfiguration (Ruff and Cohen, 2019; Bernardi et al., 2020; Martín-Sánchez et al., 2025), supporting option comparison and/or coordination across decision circuits.

We tested these possibilities in a dataset of 16,495 neurons recorded across nine frontal and subcortical areas while two macaques made multi-attribute choices (**Table S1**) (London et al., 2026). Subjects performed a one forced-choice (1FC) task were they selected a single option associated with a given reward probability (10-90%) and a juice flavor, and a two-alternative forced-choice (2AFC) task where they chose between two options differing in both attributes (**Figure 1a**). Choices in the 2AFC task were guided by the probability and a session-varying flavor preference (**Figure 1b-d**, see also (Stoll and Rudebeck, 2024a)). Eye-tracking revealed two distinct behavioral patterns in the 2AFC task. In most trials, monkeys made a single fixation toward the selected option, suggesting a quick covert evaluation of the available options (**Figure 1e**). In a smaller proportion of trials, monkeys hesitated, oscillating back and forth between options before deciding. These hesitation trials were most frequent when the options’ probabilities were similar (**Figure 1f**), suggesting that monkeys were deliberating between the options.

**Figure 1.**
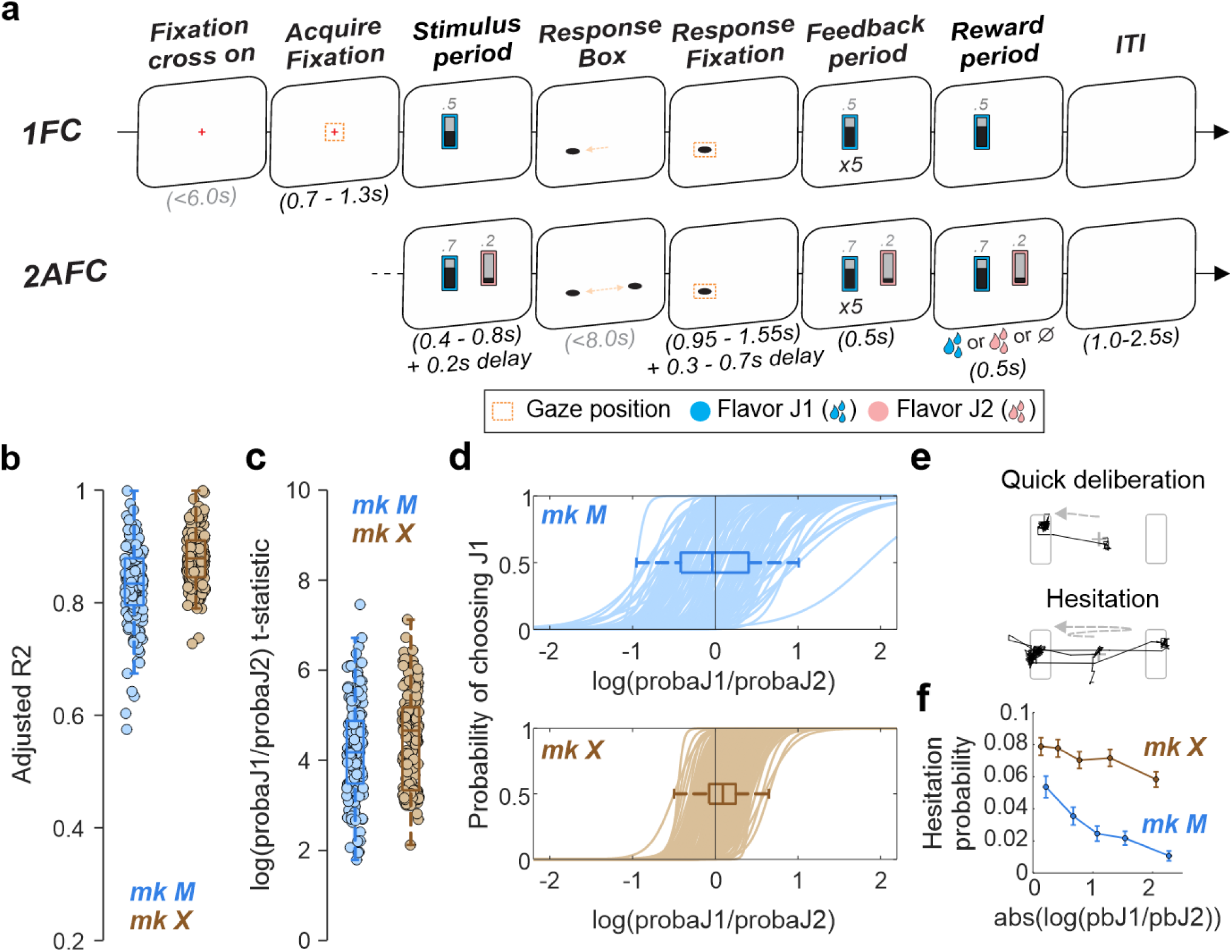
Tasks and behavioral evidence for deliberation. **(a)** Schematic representation of the behavioral tasks (1FC: top, 2AFC: bottom). Monkeys initiated trials by fixating a central fixation cross before being presented with one (1FC) or two (2AFC) stimuli. Stimuli were composed of a central bar, more or less filled, indicating the probability of being rewarded and a background color indicating the flavor of that reward in case monkeys chose that option and were ultimately rewarded. Monkeys confirmed their choice by fixating the response box on the side of the stimulus. After a delay, the chosen stimulus was flashed 5 times before monkeys received (or not) their reward given the characteristics indicated by that stimulus (flavor and probability). In 2AFC trials, the unchosen stimulus was also continuously shown on the screen during the feedback period. Trials ended after another delay (ITI: inter-trial interval). **(b)** Adjusted R² from a logistic regression predicting monkeys’ choices during 2AFC trials based on the log ratio of probabilities associated with each reward flavor (ratioPB = probaJ1/probaJ2). Dots represent individual sessions for monkeys M (mk M, blue) and X (brown), with box-and-whisker plots indicating the median (central line), interquartile range (box), and standard deviation. **(c)** Corresponding t-statistics for the log ratio of probabilities, shown as in panel b. **(d)** Predicted probability of choosing Flavor J1 against the log ratio of probabilities associated with both reward flavors for each session in monkey M (top) and X (bottom). Box-and-whisker plots indicate the flavor preference across sessions, defined as the log ratio of probabilities at which monkeys chose Flavor J1 in half of the trials. Monkeys’ choice depended on the offered probability (sigmoid function) and reward flavors (different bias from session to session). **(e)** Example eye traces during the stimulus period, with grey rectangles representing the two options and a central grey cross representing the location of the previously shown fixation cross. Quick deliberation trials (top) were characterized by a unique fixation toward one of the two options. Hesitation trials (bottom) were characterized by saccading to both sides at least once (percent of trials ± standard deviation, monkey M: 2.9±1.9%, monkey X: 7.4±4.1%; n=3928 total trials). **(f)** Probability of exhibiting hesitative behavior (mean ± 95% CI) against quintiles of absolute log ratio of probabilities centered on the flavor preference observed in each session. Monkeys were more likely to hesitate when the options presented were close to their choice indifference (mixed-effect logistic regression, F(1,67684)=99.9, p=1.6e-23).

Given this pattern of behavior, we set out to identify dynamic representational states in frontal cortex and subcortical areas during decision-making that might correspond to moments of commitment or reconsideration. We focused on reward probability as the basis for identifying these states, for three reasons. First, probability was the primary decision variable: it accounted for the largest portion of choice variance (**Figure 1b**), was encoded by a large number of neurons across areas (**Figure S1**) and could be decoded from population activity across tasks (**Figure S2a-d**). Second, probability varied across multiple levels (from 10 to 90%) which meant that decoded posteriors provided a rich, graded signal that could reveal moment-to-moment fluctuations during deliberation. Third, overall firing statistics across tasks were not only consistent (**Figure S3**) but probability representations specifically spanned a task-general neural subspace (**Figure S2e-f**). These properties allowed us to apply a cross-task decoding approach (Rich and Wallis, 2016) in which linear discriminant analysis (LDA) classifiers trained to discriminate reward probability levels from 1FC stimulus-related population responses were applied to individual 2AFC trials (**Figure 2a**). This generated trial-by-trial time-resolved posterior probabilities that tracked which option’s reward probability the neuronal population represented at each moment during the choice period.

**Figure 2.**
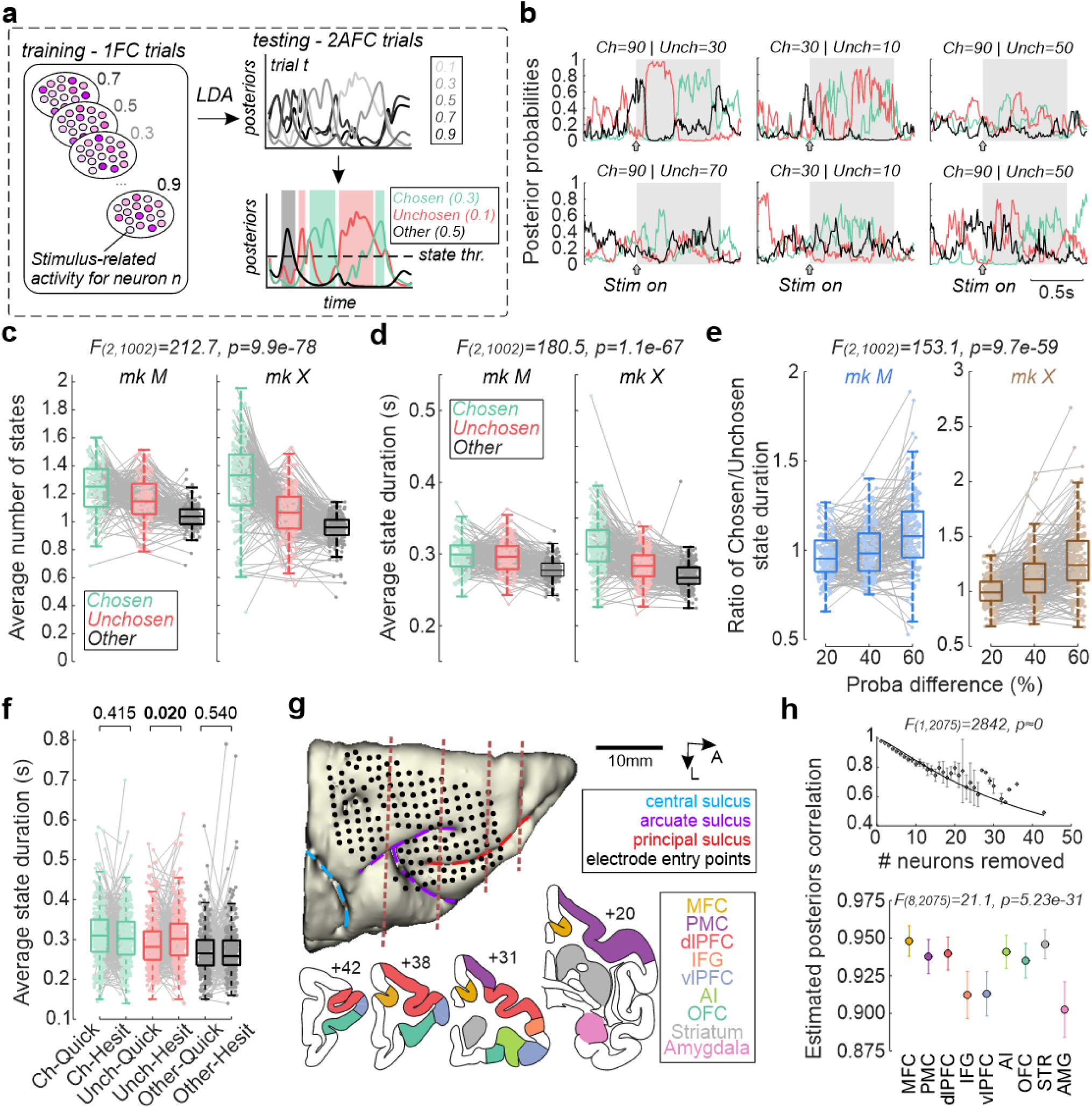
Dynamic probability states during choice. **(a)** Schematic of the cross-task decoding strategy. LDA classifiers were trained to discriminate 5 probability levels based on 1FC population responses during the stimulus period, pooling all simultaneously recorded neurons. These classifiers were then applied to 2AFC trials to generate time-resolved posterior probabilities for each probability level. For each trial, we tracked the posterior probability for the chosen option’s probability (green), the unchosen option’s probability (red), and a randomly selected probability level not offered on that trial (black). Periods in which one posterior consistently exceeded a threshold (0.3 for ≥5 consecutive bins of 100 ms) were identified as discrete probability states (shown shaded in their corresponding color). **(b)** Six example 2AFC trials (top: monkey M; bottom: monkey X) showing the temporal evolution of posteriors. Grey shading indicates the analysis window. **(c-d)** Average number (c) and duration (d) of chosen, unchosen or other probability states per trial for monkey M (left) and X (right). Chosen states (green) were significantly more frequent and lasted longer than unchosen (red) or other (black) states. Dots: sessions; grey lines connect state types within sessions. Statistics are based on a linear mixed-effects model testing the influence of state type as a fixed effect, with random intercepts included for monkeys and for sessions (nested within monkeys). **(e)** Ratio of chosen-to-unchosen state duration as a function of the absolute probability difference between options, for monkey M (blue, left) and X (brown, right). Similar statistics as panel c-d were used but now testing the influence of absolute probability difference as a fixed effect. When decisions were harder (smaller difference in probability), the chosen state was less strongly represented. **(f)** Average state duration as a function of ease of deliberation (quick or hesitation). Statistics tested the influence of state type by deliberation levels as fixed effects (interaction: F(2,1954)=2.46, p=0.08). **(g)** Recording sites. Top: Dorsal view of the 157 recording sites (black dots) targeted with the semi-chronic microdrive, overlaid on an MRI-based reconstruction of the right hemisphere of monkey X. Dashed lines represent the arcuate (violet), principal (red) and central (blue) sulci. Brown dashed lines indicate the antero-posterior levels of the coronal sections shown in the bottom panels. Bottom: Coronal sections illustrating the areas from which single neurons were recorded in monkey X. **(h)** Area contribution analysis. Posterior probability correlation between classifiers trained using all neurons or after removing neurons from a given area. Top panel: dots and error bars show the observed mean ± 95% CI of Pearson correlations across sessions and areas. The curve shows the LME model-estimated correlation as a function of neurons removed. The bottom panel shows the estimated correlation strength (± 95% CI, delta method) when removing 5 neurons (median number of neurons removed across areas and sessions) for each area. Statistics are based on a linear mixed-effects model testing the fixed effects of area and log number of neurons removed on the logit-transformed correlation strength, with random intercepts for monkeys and sessions (nested within monkeys) (conditional R²=0.68). Model estimates were back-transformed from logit to correlation (R) scale for visualization.

Posteriors from the decoder revealed clear temporal dynamics between the presented options’ probability levels during individual 2AFC trials (**Figure 2b**). We defined periods where the posterior for a given probability remained consistently above threshold (0.3 for 5 consecutive 100ms bins stepped by 10ms) as discrete probability states. Across monkeys and trials, chosen probability states were more frequent (**Figure 2c**), longer-lasting (**Figure 2d**), and occurred later (**Figure S4a**) than unchosen or ‘other’ probability states (the latter being a randomly selected probability not offered on that trial). Further, we found that the ratio of chosen to unchosen state numbers and durations were lower on trials with smaller probability difference between options (**Figure 2e** and **S4b**), indicating that state representations scaled with decision confidence. Finally, hesitation trials were characterized by longer unchosen states (**Figure 2f**) and a lower number of chosen states (**Figure S4c**) compared to quick deliberation trials, consistent with the extended evaluation of both options. Thus, the frequency, duration, and dynamics of chosen and unchosen reward probability states have a direct relationship to decision-making behavior.

We next assessed which areas contributed most to these probability states. To do this, we systematically removed each area’s neurons from the decoder and measured the impact on posteriors using correlations (**Figure 2g-h**). Three areas stood out as the strongest contributors to probability states: amygdala (AMG), inferior frontal gyrus (IFG), and ventrolateral PFC (vlPFC) (**Figure 2h**). This is consistent with these areas having the highest proportions of neuron encoding probability in both tasks (**Figure S1**) and cross-task probability decoding accuracies (**Figure S2**).

Given the distributed and dynamic nature of these probability-specific states, we next asked whether they serve a broader computational role in organizing how other option attributes are represented across frontal and limbic networks. Because each option bundles probability with other attributes (flavor identity, response side), entering a probability state associated with one option could help coordinate the representations of the other attributes across areas. To test this, we computed each neuron’s firing rate separately within each observed probability state and decoded the flavor identity or response side from pseudopopulations of increasing size (25 to 500 neurons per area; **Figures 3a** and **S5**). Critically, we trained cross-validated classifiers in either probability state and tested them in the same state or the other, producing a 2×2 matrix of decoding accuracies (train state x test state). Decoding performance was then compared across training state, testing state, pseudopopulation sizes and areas using linear mixed-effects (LME) models (see **Methods**). This cross-state decoding approach distinguishes three scenarios for how probability states could affect attribute representations (**Figure 3b**). First, probability states could scale the separability of attribute representations (state-specific gain). This alone would be expected if probability states act as an attentional spotlight, uniformly boosting the encoding of all attributes of the currently represented option across the decision network (Maunsell, 2015; Xie et al., 2018). A second alternative is that probability states could cause changes in the coding axes, regardless of potential changes in attribute separability, so that the geometry used to represent an attribute differs between states (state-specific geometry) (Bernardi et al., 2020; Chandrasekaran et al., 2025). In this case, geometry reconfiguration could be area- and attribute-specific and characterized by within-state decoders outperforming cross-state decoders. Third, attributes could be encoded along shared axes that are preserved across states (state-general geometry), in which case cross-state decoders should perform as well as within-state decoders. Such a pattern would indicate that probability states have little impact on other attributes’ representations. We distinguished between these scenarios using three summary FDR-corrected contrasts: (1) gain modulation (comparing estimated performance when tested on chosen or unchosen probability states), (2) geometry specificity (comparing within- and cross-state performance) and (3) cross-state generalization (comparing cross-state performance to chance) (see **Figure S6a** for a schematic of the contrasts). We report the following findings for pseudopopulations of 200 neurons but these patterns were robust across the tested pseudopopulation sizes (**Figure S7**)

**Figure 3.**
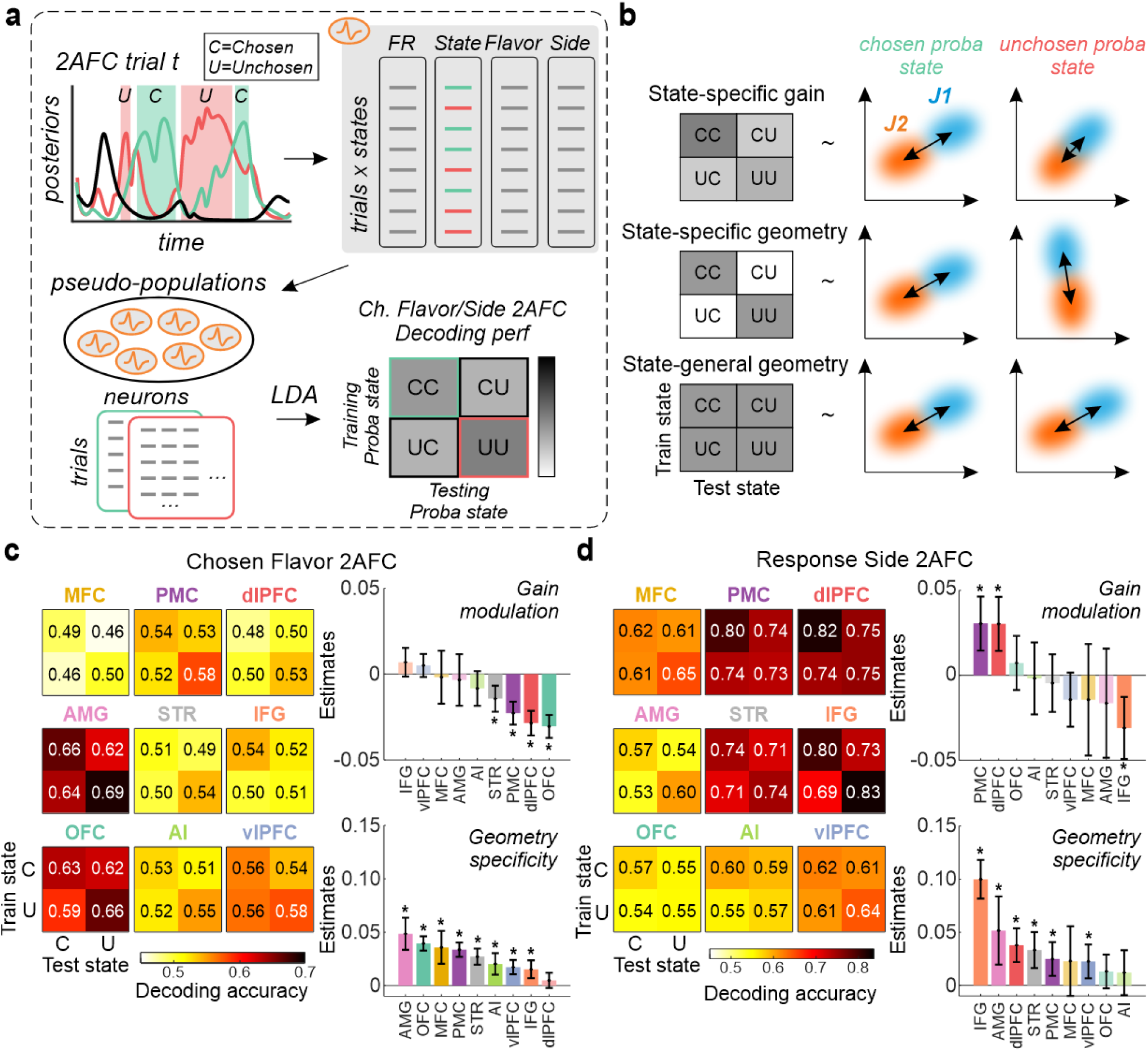
Probability states reshape population coding of other attributes in an area-specific manner. **(a)** Schematic of the cross-state decoding approach. For each neuron, firing rates were averaged within periods classified as chosen- or unchosen-probability states. Pseudopopulations of 20 to 500 neurons per area were then constructed and decoders were trained and tested within or across probability states (2×2 matrix: train-chosen test-chosen, train-chosen test-unchosen, etc.). **(b)** Illustration of the possible reconfiguration of population encoding, with 2×2 cross-decoding patterns (left column) and a representation of the underlying two-dimensional population activity across chosen and unchosen probability states (right). **(c-d)** Estimated chosen flavor (c) and response side (d) decoding accuracy across areas and probability states. Left: 2×2 cross-decoding performance estimated using a mixed-effect model with train state, test state, area and log of pseudopopulation size (and all interactions) as fixed effects (conditional R², flavor=0.83; side=0.79). Right: LME contrast estimates (± 95% CI) for the gain modulation (top) and geometry specificity (bottom) contrasts across areas. Estimated performances are shown for pseudopopulations of 200 neurons for each area. Stars: significant post-hoc differences between states (FDR-corrected).

We found that state-specific gain and geometric reshaping were most prevalent compared to state-general geometry, with probability states reconfiguring coding in an attribute- and area-specific manner (**Figure 3c-d**). For flavor, the most striking finding was a regional dissociation in the direction of state-dependent modulation across areas (**Figure 3c**). In most areas, and particularly in the orbitofrontal cortex (OFC), flavor representations were most separable during the unchosen probability state. This was not the case in areas most important for defining probability states, notably vlPFC and AMG, where no difference in flavor separability across states was observed despite strong flavor representations (see **Figure S6d** for pairwise area comparisons). Beyond this, within-state decoding exceeded cross-state decoding in most areas (**Figure 3c** and **S6b**), highlighting a genuine reconfiguration of the population geometry beyond a simple scaling change along the discrimination axis. Nevertheless, cross-state generalization remained strongest in AMG, OFC and vlPFC (**Figure S6b**), indicating that a shared component of the flavor code was preserved across states in these areas.

Representations of the upcoming response showed a related but distinct pattern (**Figure 3d**). Response side decoding was strongest in dorsal and motor-related areas (dorsolateral PFC (dlPFC), IFG, premotor cortex (PMC) and striatum (STR)), consistent with their known involvement in spatial attention and action planning (Desimone and Duncan, 1995; Goldman-Rakic, 1995; Wallis and Miller, 2003; Lebedev et al., 2004; Seo et al., 2012). Contrary to flavor, most areas decoded response side uniformly during chosen or unchosen probability states, with the exception of dlPFC showing the largest chosen-state enhancements of representations. This contrasted with the IFG population which also exhibited the strongest decoding of side but under a bias toward unchosen probability states. A few areas (most notably IFG and dlPFC) also showed geometry specificity for the response side, confirming some reconfigurations in coding axes across motor-related areas (**Figure 3d** and **S6c**). Nevertheless, cross-state generalization was substantially higher than for flavor and significant in most areas (**Figure S6c**), indicating that response side representations retain a more broadly shared coding axis across probability states. Comparing the two attributes is informative: flavor coding appears reshaped in an area-specific manner within value-related circuits and retains little cross-state structure. By contrast, coding of the response side is contained within a more uniform and shared axis across dorsal frontal and motor-related areas. This attribute-specific pattern of reshaping provides further evidence against a uniform attentional mechanism and instead points to specific population-geometric reconfigurations of information.

How do changes in population-level geometry relate to the coding patterns of individual neurons? Coding reconfiguration across probability states could arise in two ways: individual neurons could switch which option they represent across states, or instead maintain a stable selectivity while changing how strongly they contribute to the population code. To distinguish these, we identified each neuron’s preferred flavor or response side from 1FC trials and then asked, under each 2AFC probability state, whether the neuron’s selectivity aligned with the chosen or unchosen option’s attribute (**Figure S8**). Interestingly, most neurons encoded the chosen option’s attributes regardless of whether the network was in a chosen or unchosen probability state (62-75% for flavor, 61-84% for response side across areas) **(Figure S8a-b**). Only 34.9% (417/1,195) of flavor-tuned and 21.1% (418/1,981) of side-tuned neurons switched between chosen- and unchosen-flavor coding across states (**Figure S8c-d**). This indicates that population-level geometry shifts do not strictly arise from neurons switching which option they track. Instead, probability states modulate how strongly individual neurons contribute to the population code while preserving their selectivity, consistent with a re-weighting of the readout rather than a change in their tuning.

Together, our findings reveal that decision-making in primate frontal and subcortical circuits unfolds through dynamic transitions between distributed population activity states, and that these states serve as more than markers of a single attribute currently being evaluated (Rich and Wallis, 2016; Balewski et al., 2023). Here, probability states non-uniformly reshape the geometry of how other attributes are encoded in an area- and attribute-specific manner. Critically, this reconfiguration occurs despite mainly stable single-neuron selectivity, indicating that probability states shift the population coding axis for other attributes by altering which neurons most strongly contribute to attribute discriminability without changing the tuning of individual neurons (Kaufman et al., 2014; Elsayed and Cunningham, 2017; Vyas et al., 2020). Based on prior work showing attentional modulation of PFC during decision-making (McGinty et al., 2016; Xie et al., 2018), it could have been expected that probability states would homogeneously boost the coding of other attributes across areas. Instead, our results support and extend the emerging view that cognitive states reshape population geometry in a selective and area-specific way, altering which information is available for downstream read-out (Ruff and Cohen, 2019; Bernardi et al., 2020; Srinath et al., 2021). Further, the coexistence of chosen and unchosen probability states, whose balance progressively changes with decision confidence and deliberation time, supports distributed competition frameworks in which multiple options are simultaneously maintained across value and motor networks before an action is made (Cisek, 2012; Rushworth et al., 2012; Hunt et al., 2014; Thura et al., 2022; Chandrasekaran et al., 2025).

Importantly, areas that are central to generating probability states (vlPFC, IFG and AMG) show distinct state dependencies for attribute coding compared to other areas. For instance, OFC and AMG were differently influenced by probability states despite comparable flavor decoding strength, a pattern we also observed when decoding the response side using dlPFC and IFG populations. This suggests that state-generating areas might inform downstream areas which option is currently considered, with that signal shaping other attribute coding geometry. Such a mechanism is consistent with contextual gating via recurrent dynamics (Mante et al., 2013; Bernardi et al., 2020; Chandrasekaran et al., 2025; Martín-Sánchez et al., 2025) and with state-selective modulation of inter-areal communication subspaces (Semedo et al., 2019; Stoll and Rudebeck, 2024c). Taken together, our data indicate that dynamic probability states serve as distributed coordination signals that reshape how sensory and motor information are organized across the decision network during deliberation, not by creating or engaging separate coding populations for each option but by dynamically reweighting population coding axes in an area-specific manner.

## SUPPLEMENTARY MATERIALS

**Figure S1.**
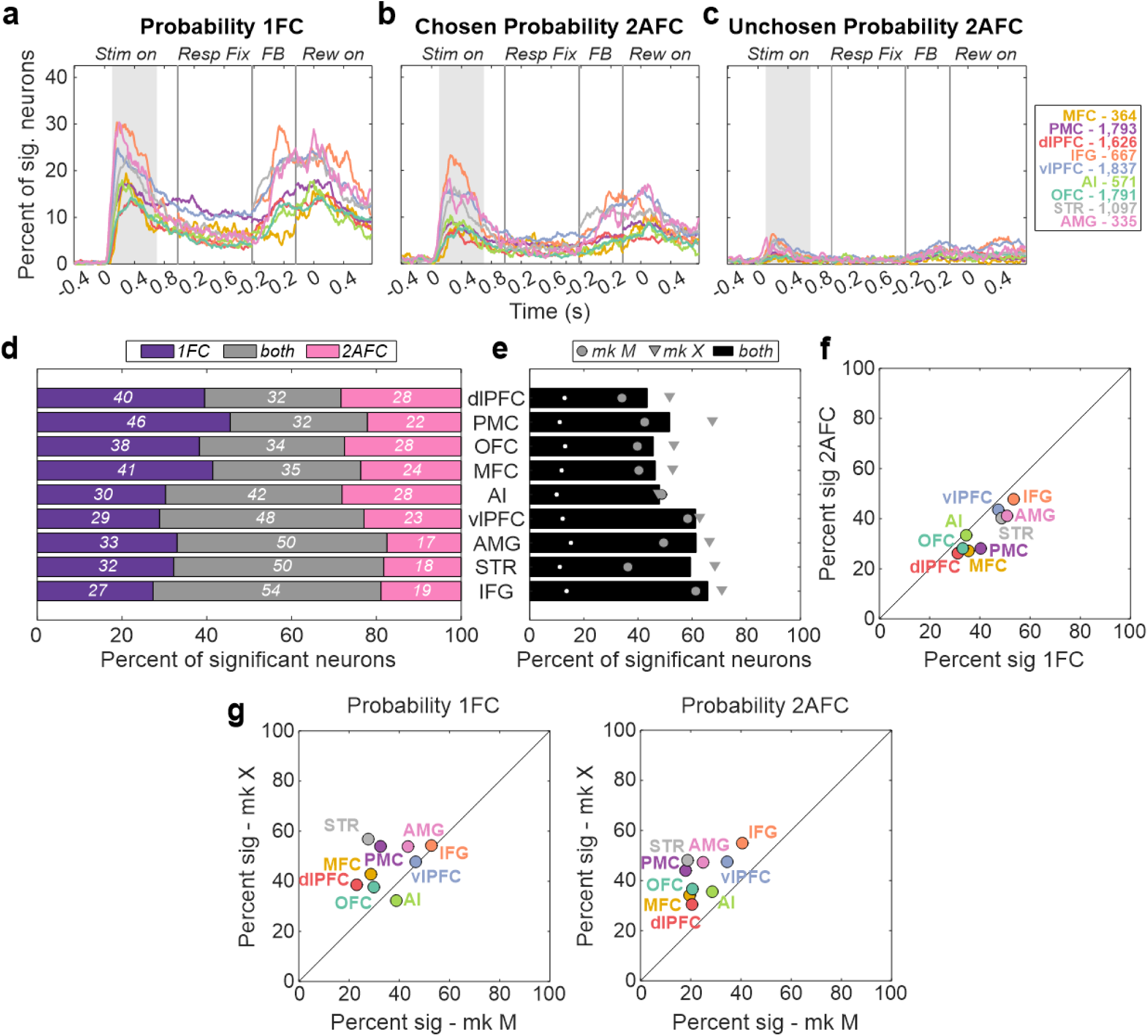
Single-neuron probability encoding across tasks. **(a-c)** Time-resolved percent of neurons per area (colors) significantly encoding (p<0.01 for ≥5 consecutive bins): (a) the stimulus probability during 1FC trials, (b) the chosen probability, and (c) the unchosen probability during 2AFC trials. Data are aligned to stimulus onset, response fixation, feedback and reward onset. Probability encoding was widespread across areas and tasks, with the highest proportions in vlPFC, IFG and AMG. **(d)** Percent of neurons encoding probability during the stimulus period in 1FC only (purple), 2AFC only (pink), or both tasks (grey). Areas are sorted by the percent encoding in both tasks. In the case of 2AFC, neurons were considered significantly encoding probability if they were tuned to either the chosen, unchosen or both probabilities. **(e)** Percent of neurons encoding probability during the stimulus period of either task, shown separately for each monkey (circles: M; triangles: X) and combined across monkeys (black bars). White dots indicate percent of significant neurons during the pre-stimulus window (-700 to -200 ms). **(f)** Percentage of significant probability encoding neurons in 1FC task vs 2AFC task. **(g)** Percentage of significant probability encoding neurons in monkey M compared to monkey X in 1FC (left) and 2AFC (right).

**Figure S2.**
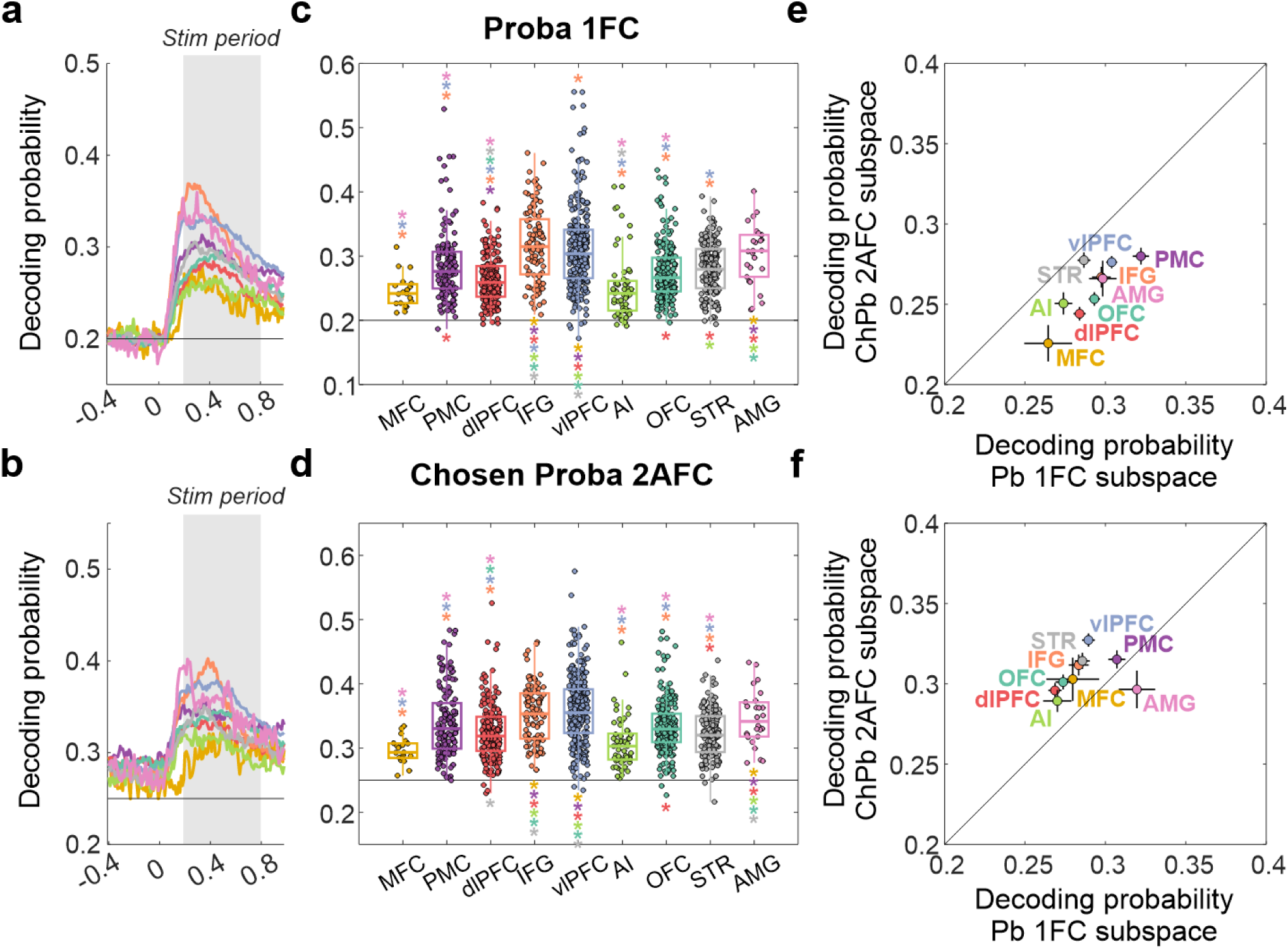
Population decoding of probability across tasks. **(a-b)** Time-resolved population decoding accuracy for reward probability in 1FC trials (a) and chosen probability in 2AFC trials (b) around stimulus onset across areas. Grey shading represents the time window used in the following panels. **(c-d)** Average stimulus period decoding accuracy for probability in 1FC (c) and chosen probability in 2AFC (d) trials across sessions (dots) and area (colors). Statistical significance was assessed using generalized mixed-effect models, with area and number of neurons as factors and monkey/session as random intercepts (factor area: probability 1FC, F(8,1148)=34.9, p=1.1e-49; chosen probability 2AFC, F(8,1147)=35.3, p=4.2e-50). Star’s locations (FDR-corrected p < 0.01) indicate the direction of the effect (e.g., yellow star, MFC, under the orange boxplot, IFG, means that MFC decoding performance was significantly worse than IFG). The horizontal bar represents the theoretical chance level. **(e-f)** Average decoding accuracy for probability in 1FC (e) and chosen probability in 2AFC (f) after projecting the population activity onto a probability subspace defined on 1FC trials (x-axis) against the one defined on 2AFC trials (y-axis). Decoding accuracies were above chance level for all areas and regardless of which probability subspace the population activity was projected onto, revealing that probability representations occupy a task-general neural subspace.

**Figure S3.**
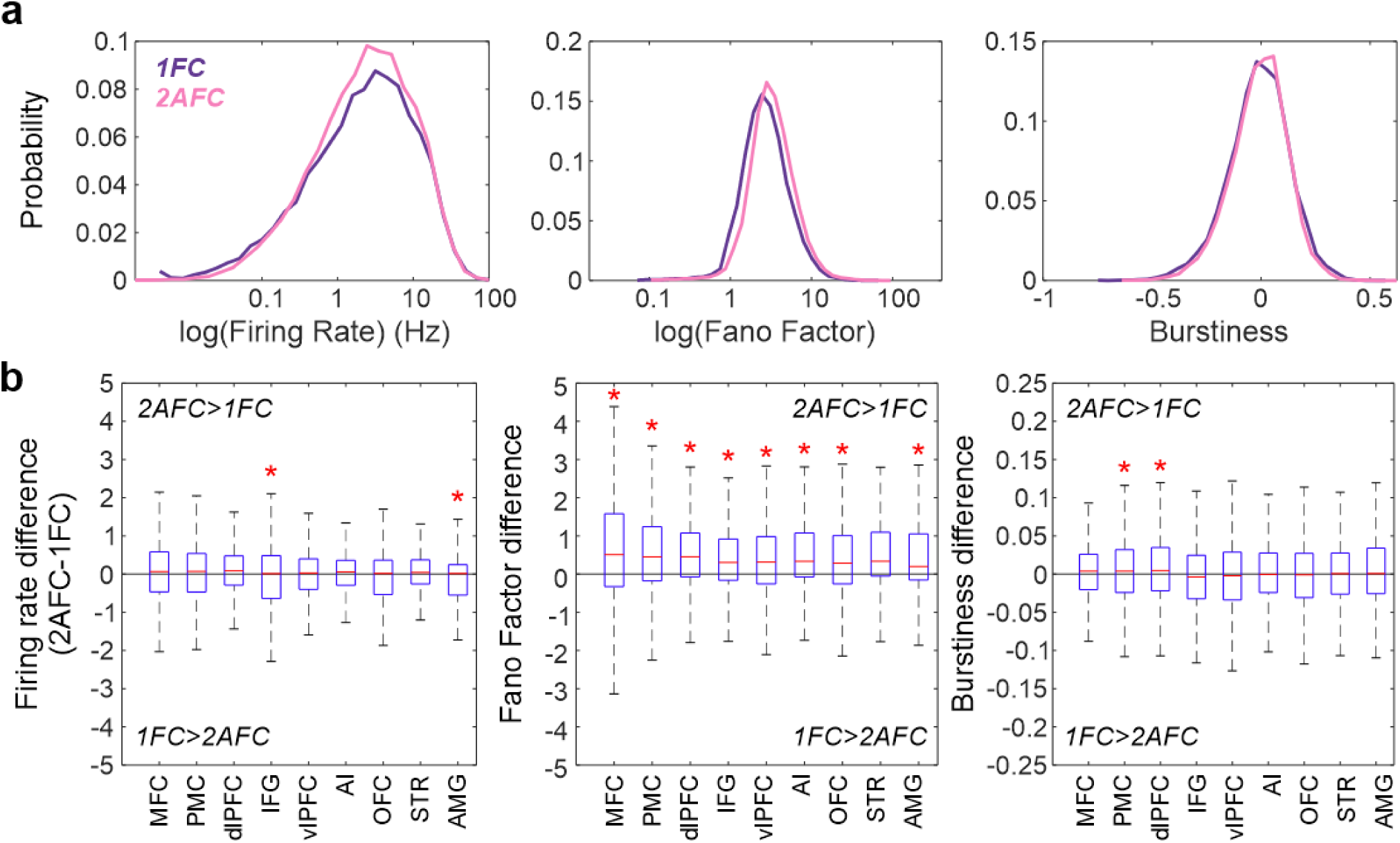
Spiking statistics across tasks and areas. **(a)** Distributions of log mean firing rate (left), log Fano factor (middle), and burstiness index (right) during the 1FC (purple) and 2AFC (pink) tasks. **(b)** Boxplots show the distribution of per-neuron difference in firing rate (left), Fano factor (middle) and burstiness (right) between 2AFC and 1FC tasks across areas. Statistical significance was assessed using LME models for each area with monkey as a random intercept (diff ∼ 1 + (1|monkey)), testing whether the mean difference was significantly different from zero (stars indicate p<0.01). Spiking statistics of neurons remained comparable across tasks, except for greater Fano factor in 2AFC trials in neurons from most areas.

**Figure S4.**
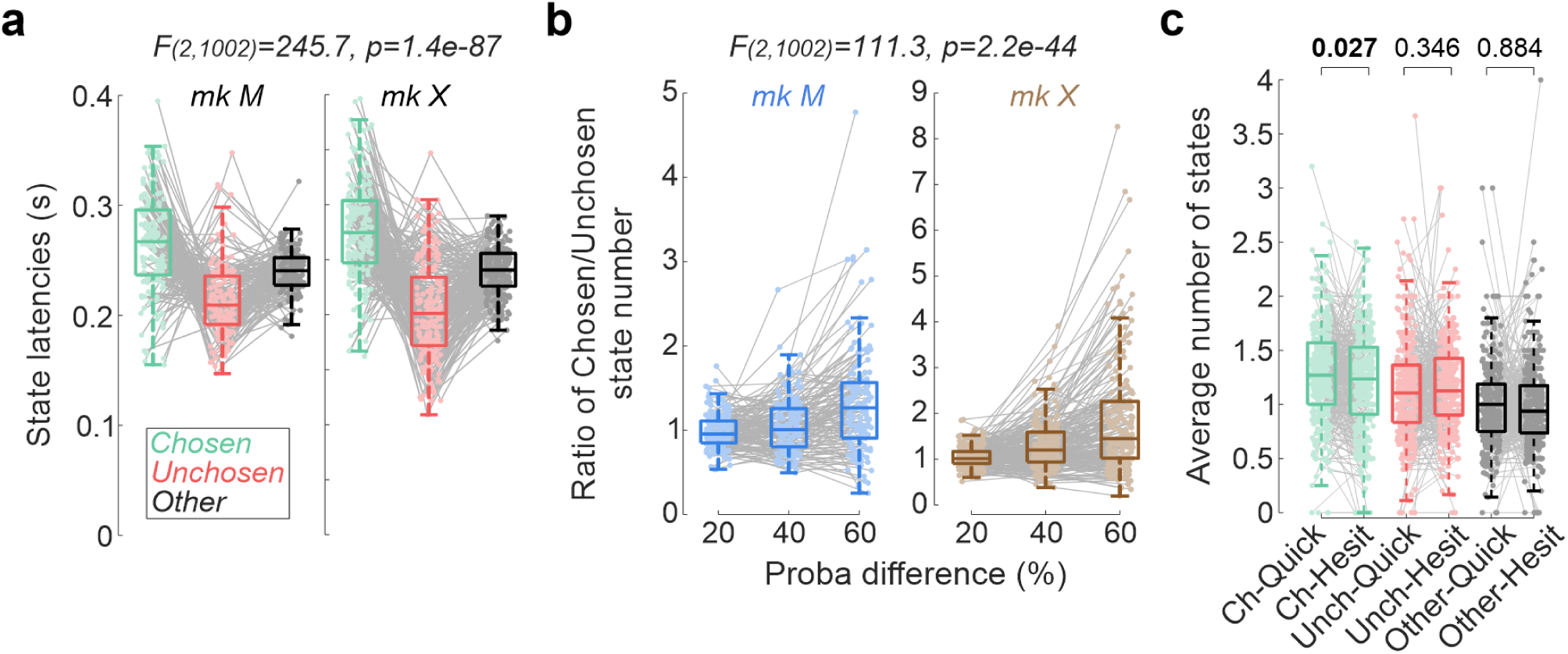
Probability state dynamics: additional statistics. **(a)** Average latency of chosen (green), unchosen (red), and other (black) states relative to stimulus onset. Conventions as in Figure 2c-d. Unchosen states tended to emerge earlier than chosen states. **(b)** Chosen-to-unchosen ratios for number of states as a function of the absolute probability difference between options. As decisions became easier (large probability difference), the chosen-state advantage increased. **(c)** Average number of states as a function of ease of deliberation (quick or hesitation) and state type (colors). Quick deliberation trials were characterized by an increased number of chosen states (interaction: F(2,1980)=2.58, p=0.076).

**Figure S5.**
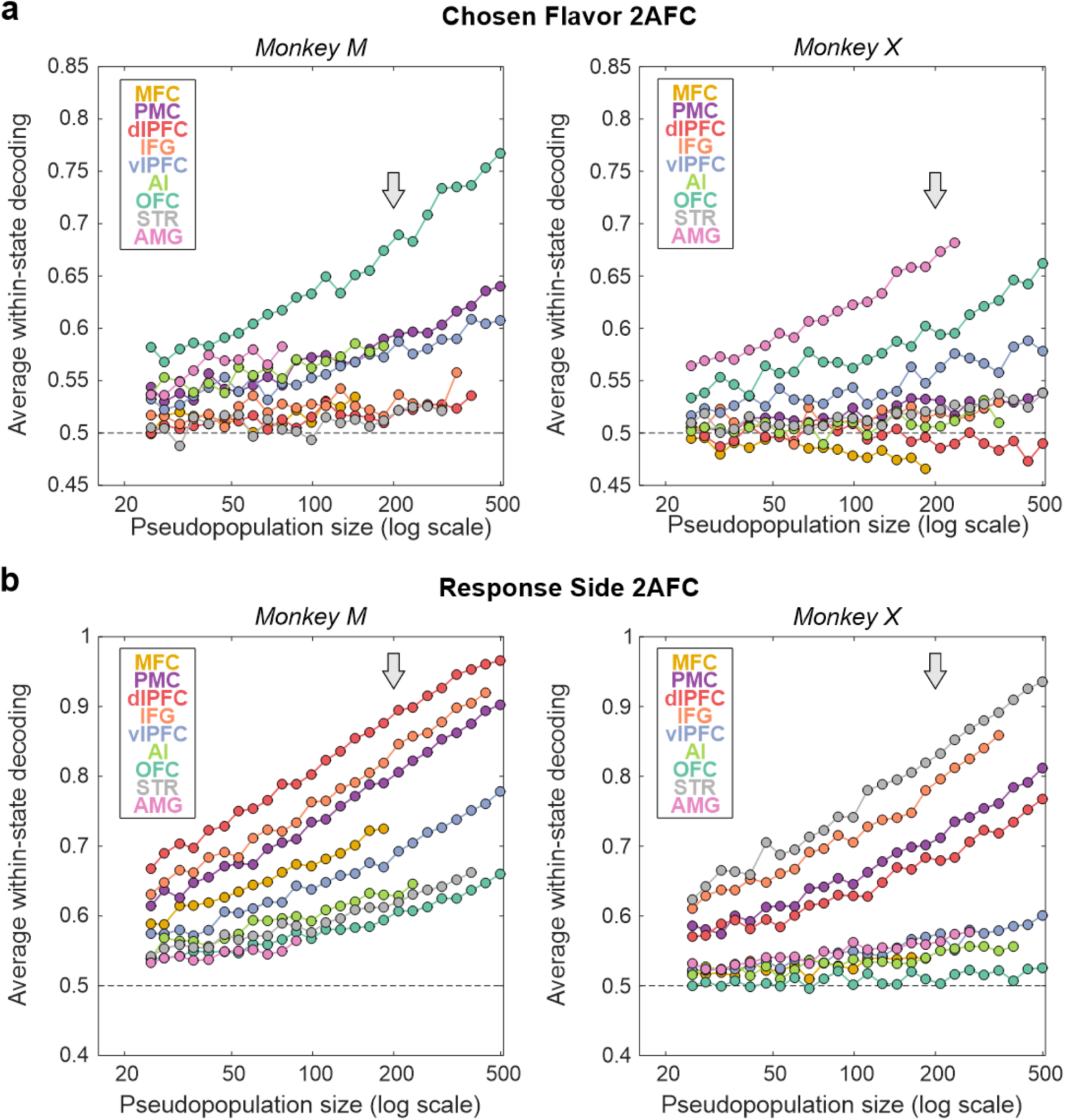
Population-size curves for chosen flavor and response side decoding across states. **(a-b)** Average within-state decoding accuracies for chosen flavor (a) and response side (b) as a function of pseudo-population size (25-500 neurons, log scale) for each area and monkeys (left: monkey M, right: monkey X). Curves show the mean performance across 100 pseudo-population draws. Grey arrows represent the pseudo-population size at which contrasts are reported in the main text (see also **Figure S7**).

**Figure S6.**
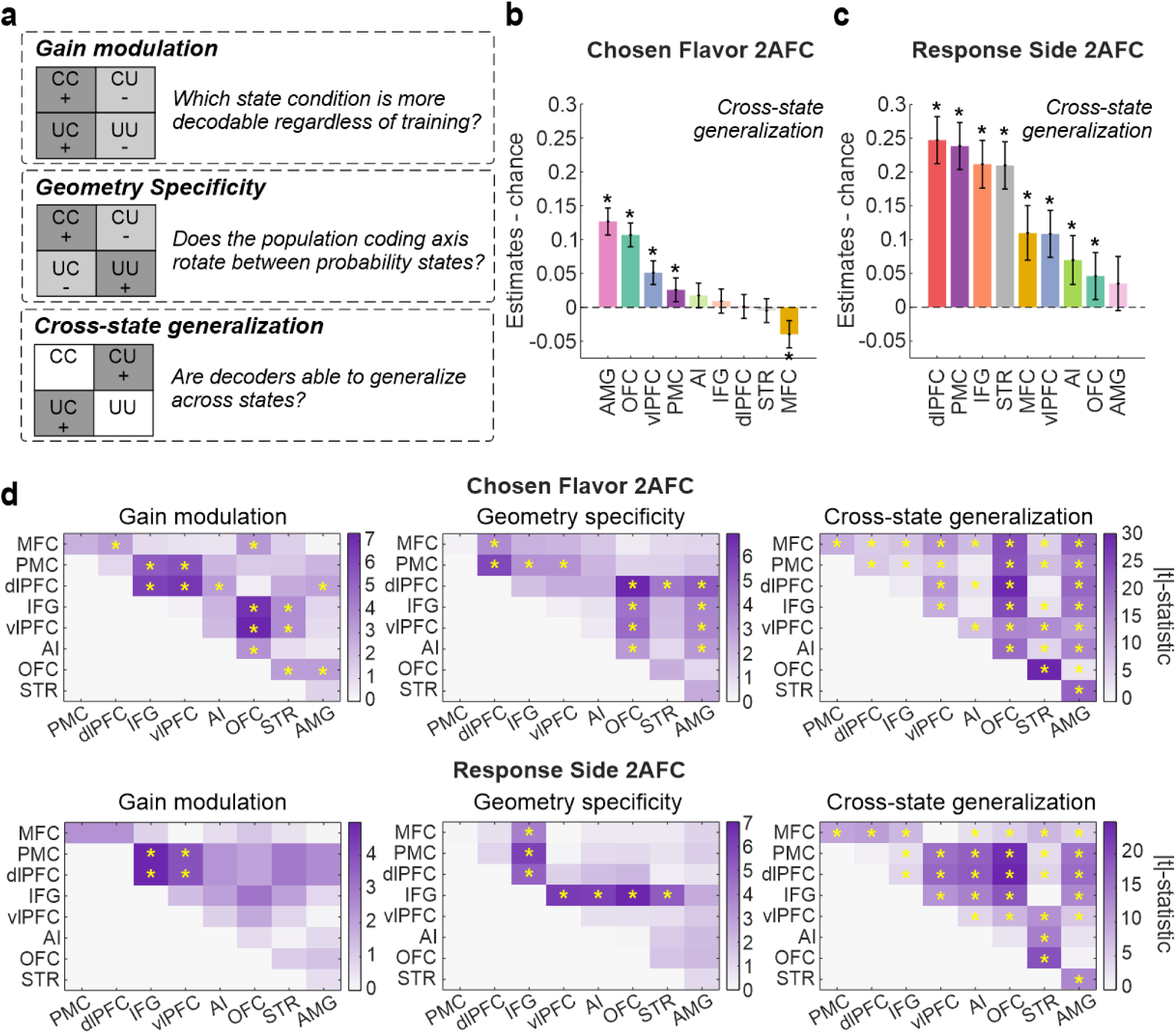
Definitions and estimates of cross-state decoding contrasts. **(a)** Schematic illustrating the three post-hoc contrasts derived from the 2×2 train by test state decoding matrix (train-test: C-C, C-U, U-C, U-U, where C=chosen, U=unchosen). Gain modulation: tested-on-chosen performance minus tested-on-unchosen performance; Geometry specificity: within-state performance minus cross-state performance; Cross-state generalization: cross-state performance (off-diagonal) minus chance. **(b-c)** Estimates (± 95% CI) for the cross-state generalization contrast for chosen flavor 2AFC (b) and response side (c). As for Figure 3c-d, effect sizes are shown for a pseudopopulation of 200 neurons for each area, with stars highlighting areas with significant post-hoc differences (FDR-corrected at p<0.05). **(d)** Absolute Wald t-statistic (color) for pairwise area comparisons based on the LME-estimated contrasts (geometry specificity, gain modulation, and cross-state generalization) for chosen flavor (top) and response side (bottom). Stars indicate significant pairwise area differences (FDR-corrected p<0.01).

**Figure S7.**
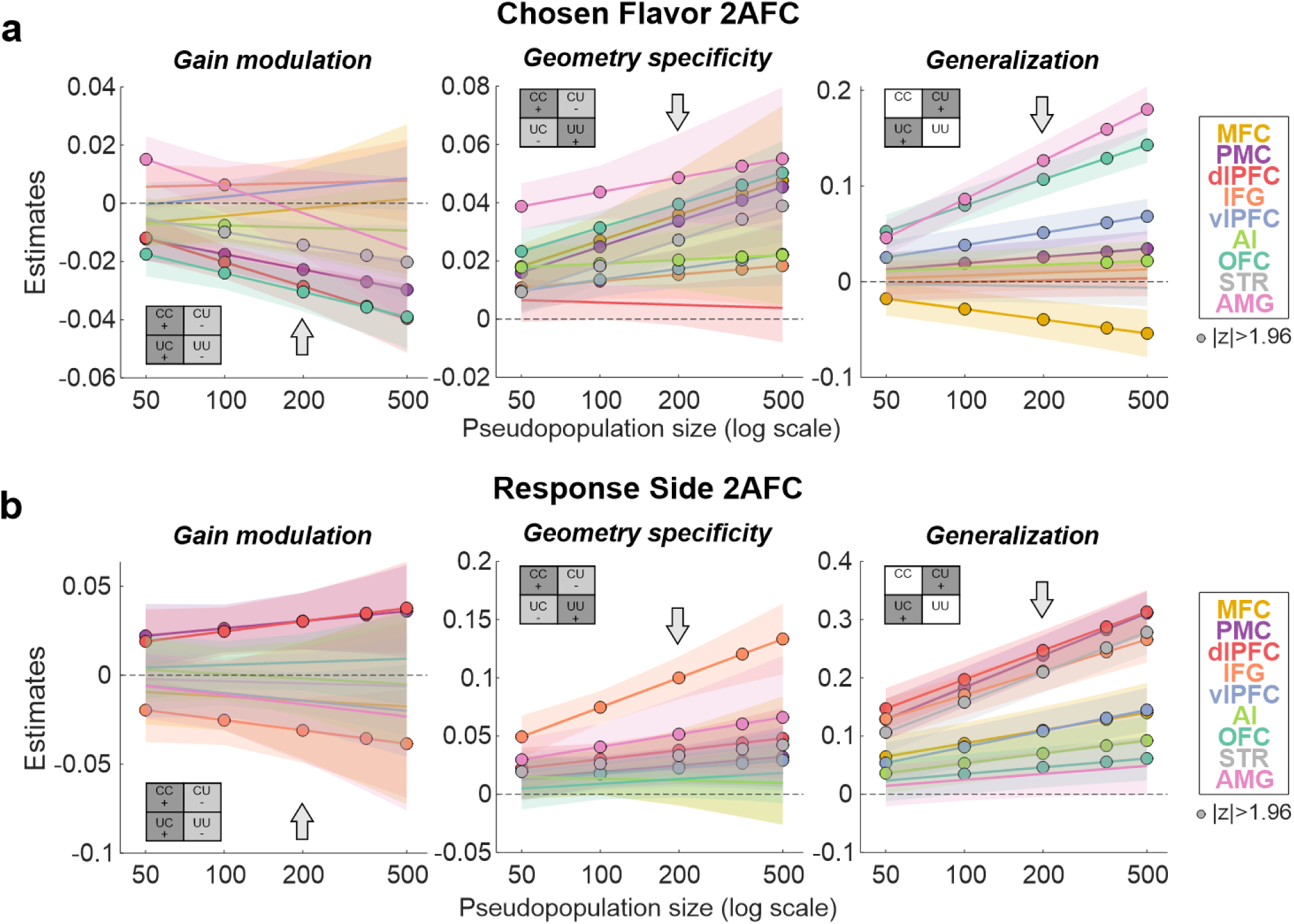
Robustness of decoding contrasts across population sizes. **(a-b)** Post hoc estimates (± 95% CI) from LME for gain modulation (left), geometry specificity (middle) and cross-state generalization (right) contrasts for chosen flavor (a) and response side (b) for each area and as a function of pseudo-population sizes (log scale). Inset 2×2 matrices illustrate the contrast considered. Dots represent significant estimates (|z|>1.96). Grey arrows represent the pseudo-population size at which contrasts are reported in the main text. The direction (sign) and significance of the reported effects for all considered contrasts are consistent across pseudopopulation sizes.

**Figure S8.**
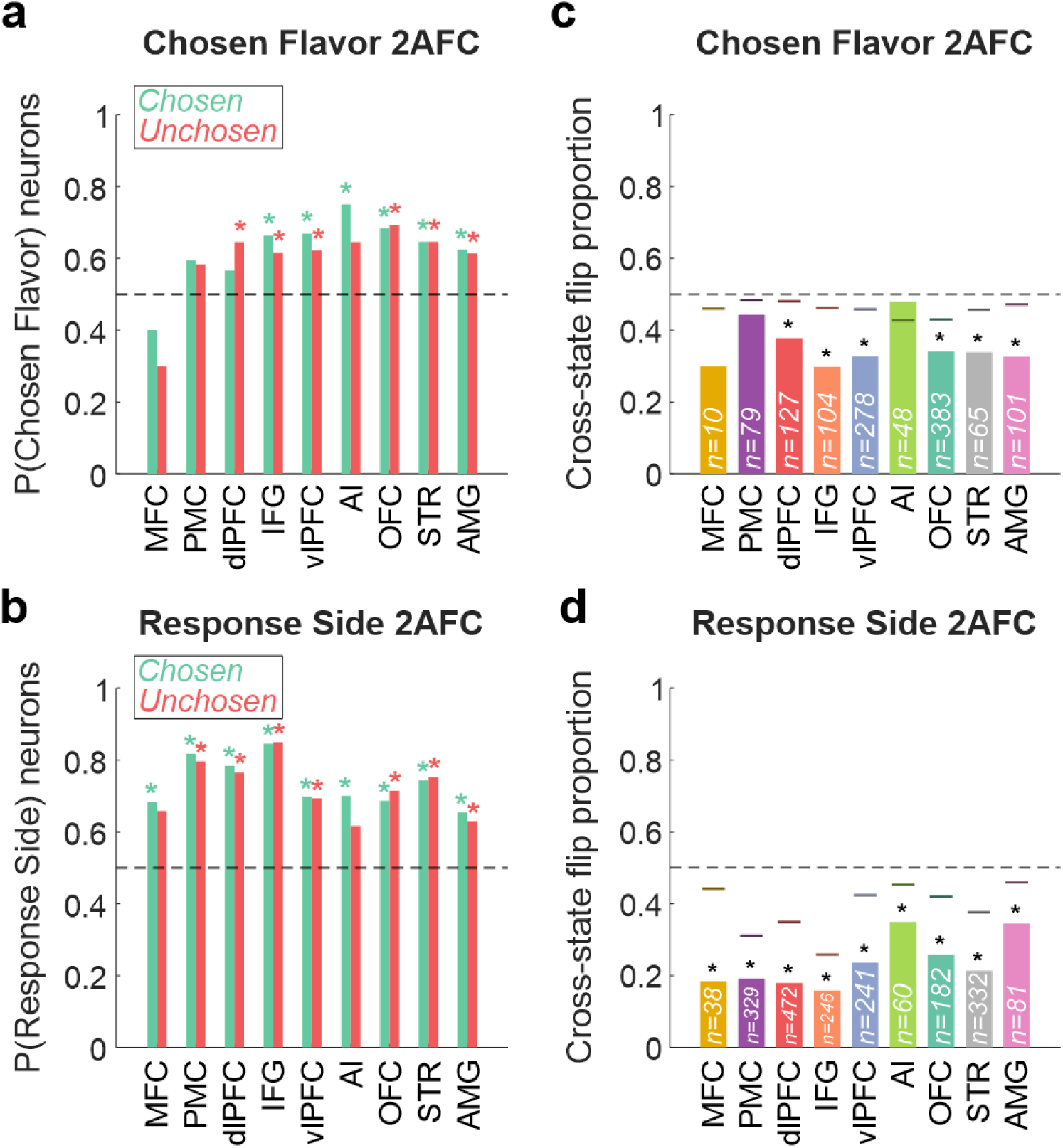
Single-neuron encoding across probability states. **(a)** Proportion of neurons encoding the chosen flavor during chosen (green) and unchosen (red) probability states across areas. Stars represent a significant deviation from chance (two-sided binomial test, FDR-corrected across areas). **(b)** As in panel (a) but for the response side. **(c)** Proportion of neurons flipping their encoding of chosen or unchosen flavor across states. Horizontal bars represent the expected proportion under independence assumption. Stars represent significant differences compared to 50% chance using FDR-corrected two-sided binomial tests. **(d)** As in panel (c) but for the response side.

**Supplementary Table S1.** Neuron counts across monkeys and areas.

| Area | Monkey M | Monkey X | Total |
| --- | --- | --- | --- |
| **MFC** | 281 | 231 | **512** |
| **PMC** | 1,894 | 1,013 | **2,907** |
| **dIPFC** | 1,362 | 1,256 | **2,618** |
| **IFG** | 599 | 496 | **1,095** |
| **vIPFC** | 1,211 | 1,909 | **3,120** |
| **AI** | 355 | 552 | **907** |
| **OFC** | 1,664 | 1,173 | **2,837** |
| **STR** | 640 | 1,234 | **1,874** |
| **AMG** | 162 | 463 | **625** |
| **Total** | **8,168** | **8,327** | **16,495** |

## METHODS

### Subjects and apparatus

Two adult rhesus macaques (Macaca mulatta), monkeys M and X, participated in the experiments. At the onset of recordings, the animals were 8 and 5.5 years old and weighed 11.9 and 7.9 kg, respectively. They were group-housed under a 12-h light/dark cycle with unrestricted food access; water was regulated five days per week during training and experiments. All procedures were approved by the Icahn School of Medicine Institutional Animal Care and Use Committee.

During recordings, animals were head-fixed 56 cm from a 19-inch monitor. Gaze was tracked at 90 Hz (PC-60, Arrington Research). Behavioral control used the MonkeyLogic toolbox in MATLAB (R2014b) and fluid rewards were delivered through a custom air-pressure–driven system (Mitz, 2005). Raw electrophysiology was acquired at 40 kHz (OmniPlex, Plexon) and spike-sorted offline using the MountainSort plugin of MountainLab (Chung et al., 2017). Only well-isolated neurons were further processed (isolation > 0.75, noise overlap < 0.2, SNR > 0.5, firing rate > 0.05 Hz). A detailed description of this dataset can be found in London et al (London et al., 2026).

### Behavioral task

During each session, monkeys performed a one-alternative forced choice (1FC) and a two-alternative forced choice (2AFC) task. In the 1FC task, a single stimulus was presented on either side of the screen and monkeys selected it by fixating the corresponding response box to earn a fluid reward. The 2AFC task required monkeys to evaluate and choose between two stimuli displayed simultaneously on opposite sides of the screen. Across both tasks, stimuli are composed of an outer colored rectangle signaling the juice flavor (out of 2 possible flavors) monkey can earn on that trial and a centrally positioned more or less filled bar whose extent indicates the probability of receiving a reward. Each session drew from two colors (chosen from a set of nine) mapped to two juice flavors (randomly selected from five: apple, cranberry, grape, pineapple, or orange; diluted to 50% in water). Reward probabilities spanned 10-90%, sampled in steps of 20% for monkey M and 10% for monkey X.

Trials began with fixation of a central cross (0.7-1.3 s, in 0.3 s steps), after which one or two stimuli appeared on the left and/or right side of the screen for 0.4-0.8 s (0.2 s steps, pseudorandomly). Following a 0.2 s blank screen, one or two response boxes appeared on the side of the previously displayed stimuli (at one of three equidistant locations: bottom, center, or top). Monkeys were then able to indicate their choice by fixating the response box next to the preferred stimulus within 8 s and required to hold fixation there for at least 0.25 s to register a response at which point the unchosen response box disappeared (in the 2AFC trials). This dwell-time requirement allowed monkeys to change their mind before committing to a choice. Monkeys then maintained fixation on the chosen response box for an additional 0.6-1.2 s (0.3 s steps). This was followed by a blank screen for a 0.3-0.7 s delay (0.2 s steps). During the ensuing feedback period, both options (or the only available one during 1FC trials) reappeared at their original locations with the chosen option flashing five times (0.1 s on / 0.1 s off) before remaining on screen for the duration of reward delivery and an additional 0.5 s. On rewarded trials, monkeys received 2-3 fluid pulses of 0.03-0.06 s each (separated by 0.1 s; 0.25-0.36 mL total) of the flavor and at the probability associated with the chosen option. Unrewarded trials were matched in duration to rewarded ones. Rewarded trials were followed by a 2 s intertrial interval and unrewarded trials by 3.5-4 s. Fixation breaks or failure to initiate a trial within 6 s triggered a 1 s red circle at screen center and an extended intertrial interval (4-6 s for monkey M; 3-4 s for monkey X).

In 2AFC trials, the two options differed in juice flavor (different-flavor trials) in 50% of trials for monkey M and 75% for monkey X. For monkey M, different-flavor trials always paired options with distinct probabilities; for monkey X, probabilities could be identical or different across the two options. The remaining trials (50% for monkey M; 25% for monkey X) presented options of the same flavor at different probabilities (same-flavor trials).

### Choice behavior

Monkeys’ choices were analyzed using logistic regression (*fitglm* in MATLAB), restricted to trials in which the two simultaneously offered options were associated with different flavors of juice. For each session, we fitted the following model:

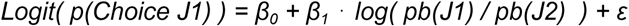

where Logit(p)=log_e_(p / 1-p), pb(J1) and pb(J2) are the reward probabilities associated with flavors J1 and J2, respectively, β_x_ are the coefficient estimates, and ε the residuals. The model used a binomial distribution with a logit link function and a Jeffreys prior penalty to stabilize estimates in sessions with limited data. We extracted the t-statistic and estimate for log(pb(J1) / pb(J2)) and the adjusted R² as measures of overall probability sensitivity per session. We also derived a session-level flavor bias (indifference point) as the value of log(pb(J1) / pb(J2)) at which predicted p(Choice J1)=0.5, computed as -β_0_ / β_1_. Exponentiating the indifference point yields an indifference ratio (pb(J1)∶pb(J2)), where a ratio<1 indicates a bias toward J1 and a ratio>1 indicates a bias toward J2. Predicted choice probabilities were computed over a continuous range of log(pb(J1) / pb(J2)) corresponding to the 10-90% probability range (*predict* in MATLAB).

### Saccade detection and trial classification

Eye movements were analyzed to determine whether animals looked directly at one stimulus or shifted gaze between both stimuli during deliberation. For each session, a fixation baseline was established by pooling all gaze samples from the fixation cross onset to the stimulus onset across trials. The fixation center was defined as the median of the horizontal and vertical gaze position across these samples, and the fixation radius was set to ensure that 55 percent of the gaze points fell within it. This threshold was established empirically to account for noise in gaze data. Saccades were then detected within the stimulus presentation window on a trial-by-trial basis, defined as horizontal excursions beyond the fixation radius lasting at least 200 ms. Saccades’ direction was determined using the gaze position relative to the fixation center (rightward or leftward). Consecutive excursions to the same side were merged into a single saccade event. For trials with no excursion meeting the duration threshold, a single saccade was assigned at the time of first fixation zone departure. For each trial, the total number of saccades, their onset times and directions, the latency of first fixation zone departure, and the entry and exit times for each crossing were extracted relative to stimulus onset. Trials with a saccade toward a unique direction were classified as quick deliberation while trials with saccades toward both sides were classified as hesitations.

### Neural recordings

Across a series of aseptic surgeries under anesthesia, monkeys were implanted with a titanium headpost and a PEEK chamber housing a 157-channel semi-chronic microdrive (Gray Matter Research) with glass-coated electrodes (1-2 MΩ at 1 kHz; Alpha Omega). Electrodes were regularly moved to record from distinct neurons and areas over many months. Recording locations were confirmed via daily electrode depth monitoring, CT scans registered to pre-op and post-op MRIs, as well as post-mortem histology. Detailed description of the histological confirmation and identification of the areas was previously published (London et al., 2026). A total of 16,495 single neurons were recorded in 9 areas across 340 sessions: medial frontal cortex (MFC: area 24c, n=512), premotor cortex (PMC: area 6DR, 6DC and 6Va/Vb, n=2,907), dorsolateral prefrontal cortex (dlPFC: area 8A, 8B, 46v, 46df and 46d, n=2,618), inferior frontal gyrus (IFG: area 44 and 45, n=1,095), ventrolateral prefrontal cortex (vlPFC: area 12r, 12m, 12l and 12o, n=3,120), agranular insula (AI: n=907), orbitofrontal cortex (OFC: area 11m/l, 13m and 13l, n=2,837), striatum (STR: caudate and putamen, n=1,874), and amygdala (AMG: n=625) (**Table S1**).

To confirm that spiking statistics were comparable across tasks, we computed three measures for each neuron aligned to stimulus onset (-750ms to +1250ms). Mean firing rate was estimated as the average spike density across all trials and time bins within this time period. The Fano factor was computed as the ratio of the trial-to-trial variance to the mean of the total spike count per trial. The burstiness was quantified as the normalized coefficient of variation of inter-spike intervals (ISIs) following the equation:

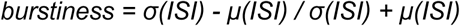

and computed per trial from trials with more than 5 spikes. The difference between 2AFC and 1FC for these three measures is reported on **Figure S3**.

### Single neuron encoding of decision-related variables

Single-neuron encoding was assessed separately for each task using multi-way ANOVAs applied at each time bin to min-max normalized firing rates (**Figure S1**). For 1FC trials, a three-way ANOVA was fit with reward probability (5 levels: 10/30/50/70/90%, treated as a linear covariate), juice flavor (categorical, 2 levels), and response side (categorical, 2 levels) as factors. For 2AFC trials, a four-way ANOVA was fit with chosen reward probability (4 levels: 30/50/70/90%, linear), unchosen reward probability (4 levels: 10/30/50/70%, linear), chosen flavor (categorical, 2 levels), and response side (categorical, 2 levels). No interaction terms were included in either model. To account for trial number differences across tasks, 2AFC trials were subsampled condition-by-condition to match the number of available 1FC trials. A neuron was classified as significantly encoding a factor if the ANOVA p-value was below 0.01 for at least 5 consecutive time bins within the 200-700 ms post-stimulus window. The proportion of firing-rate variance explained by each factor and time bin was quantified using omega-squared.

### Within-task probability decoding and subspace alignment

To verify that reward probability was reliably encoded in population activity within each task, and to establish that 1FC and 2AFC probability representations share a common neural subspace, we performed two analyses using the within-session populations from each area. Sessions with fewer than 5 neurons in a given area and neurons with a mean firing rate lower than 0.5 Hz (averaged across all trials and time bins from both tasks combined) were excluded.

For each area, time bin, and task separately, a diagonal-covariance linear discriminant analysis (LDA) was trained and tested using 10-fold cross-validation to discriminate reward probability levels from single-neuron firing rates (probability during 1FC trials: 5 levels 10-90%; chosen probability during 2AFC trials: 4 levels 30-90%, unchosen probability during 2AFC trials: 4 levels 10-70%) (**Figure S2a-d**). Decoding accuracy was averaged across folds. The average performance over the 200-700 ms stimulus window was used as the session-level summary statistic and compared across areas using a generalized linear mixed effect (LME) model with area as a fixed factor and monkey and session as random intercepts.

To test whether probability representations occupy a task-general neural subspace, we extracted a one-dimensional probability coding axis from each task’s population response and assessed whether activity projected onto that axis could be decoded regardless of which task defined it (**Figure S2e-f**). For each task, the mean firing rate per probability condition during the stimulus period (200-700 ms) was computed and z-scored across conditions per neuron. We then used PCA to extract the first principal component, which defined that task’s probability subspace. Individual trials from either task were then projected onto this axis (using the z-score parameters from the condition averages), and another diagonal-covariance LDA with 10-fold cross-validation was used to decode probability from the projected values. This was done for all combinations of projection subspace (1FC or 2AFC) and decoded variables (probability during 1FC trials, chosen probability during 2AFC trials).

### Cross-task population decoding of probability

As previously, neurons with a mean firing rate < 0.5 Hz were excluded. For each session, firing rates were min-max normalized separately for each task, using each neuron’s minimum and maximum firing rate within a pre-stimulus baseline window (-900 to -100 ms) as the scaling bounds. A diagonal-covariance LDA classifier was then trained on the average normalized firing rate of neurons across all areas during the 1FC stimulus period (200-700 ms following stimulus onset) to discriminate the five reward probability levels (10-90%). The trained classifier was then applied to individual 2AFC trials using a sliding window (100 ms bins with 10 ms steps) to generate time-resolved posterior probabilities for the chosen probability level, the unchosen probability level, and a randomly drawn unoffered probability level (referred to as “other” state) (**Figure 2a**). A probability state was defined as a contiguous period in which the posterior probability exceeded 0.3 and remained the highest of the three possible states for at least 5 consecutive bins. State identity, count, duration (when a state was detected), and onset latency were extracted for each trial. State-related statistics were modeled using LME models including state type and/or decision difficulty as fixed effects and random intercepts for monkey and session.

### Neuron-dropping analysis

To quantify the contribution of each brain area to the cross-task probability representation, neurons from one of the 9 areas were removed from both the 1FC training population and the 2AFC test population, and the diagonal-covariance LDA was re-run on the reduced neuron set. We then computed the Pearson correlation of the chosen and unchosen probability posteriors obtained in the full-population compared to the area-removed ones for each time and trials. A lower correlation therefore indicates a greater contribution from the removed area. This was repeated for each area and each session. Correlations were modeled with a LME model with area (9 levels, categorical), the log-transformed number of neurons removed, and their interaction as fixed effects, and with random intercepts for monkey and session. Model estimates were extracted at the median number of removed neurons (n=5) and back-transformed from the logit scale (**Figure 2h**).

### Cross-state pseudopopulation decoding of chosen flavor and response side

For each area, pseudopopulations were built for each monkey by pooling neurons across sessions with independent trial shuffling (**Figure 3a**). Only neurons recorded for at least 20 different-flavor trials per condition were included in the following analyses. To remove possible behavioral confounds, we only used the firing rate of neurons during trials where both chosen and unchosen probability states were detected. To account for the different number of recorded neurons across areas, we created multiple pseudopopulation activity matrices with sizes ranging from 25 to 500 neurons (25 log-spaced steps, **Figure S5**), each one being randomly sampled 100 times. Decoding of chosen flavor and response side was then evaluated across all four train by test probability-state combinations (train-chosen and test-chosen, train-chosen and test-unchosen, train-unchosen and test-chosen, train-unchosen and test-unchosen) using diagonal-covariance LDA (with 10-fold cross-validation). To prevent near-silent neurons from dominating the decoder, a scaled variance floor (1% of the mean pooled variance across neurons) was added to each neuron’s pooled variance before computing discriminant weights, equivalent to L2 ridge regularization with a data-adaptive penalty. Decision scores from all cross-validation folds were normalized within each fold by the training-class separation and pooled before classification so that the training-state midpoint served as a common threshold. Decoding accuracies were then averaged across the 100 random pseudo-population sampling and entered into the following unified LME model:

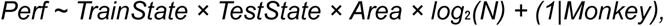

where N is the population size. Models were primarily evaluated at N=200 neurons (log₂(N)=7.64) but comparable results were observed at other population sizes (**Figure S5** and **S7**). Three orthogonal summary contrasts were derived from model estimates: (i) gain modulation, comparing estimated decoding performance when tested on chosen versus unchosen probability states; (ii) geometry specificity, comparing within-state to cross-state decoding performance; and (iii) cross-state generalization, comparing cross-state decoding performance to chance (**Figure S6a**). All contrasts were FDR-corrected across the 9 areas.

### Single-neuron tuning during probability states

#### Sign-consistency

To assess whether individual neurons encoded the chosen or unchosen flavor during each probability state (**Figure S8**), we first extracted the regression coefficient relating neurons’ firing rate to flavor identity for each significantly tuned neuron in the 1FC task (p<0.01 for ≥5 consecutive bins within 200-700 ms following stimulus onset). The sign of this coefficient defined the neuron’s preferred flavor direction. We then computed the mean firing rate across all time bins assigned to each probability state during 2AFC trials and regressed this state-average activity on the chosen flavor identity. The analysis was restricted to trials where chosen and unchosen flavors differed, making them orthogonal predictors. A neuron was then classified as encoding the chosen flavor if its 2AFC regression coefficient had the same sign as its 1FC coefficient, and unchosen flavor if signs differed. We reported the probability of encoding the chosen flavor for each area and probability state and tested deviation from chance with FDR-corrected two-sided binomial tests. A similar analysis was conducted to assess response side tuning.

#### Cross-state flip score

For each neuron, we also compared its chosen/unchosen flavor tuning across the two probability states. A neuron is said to “flip” if it encoded the chosen flavor in one state and the unchosen flavor in the other. The flip rate per area was computed for both flavor and response side and tested against 50% chance (two-sided binomial test, FDR-corrected across areas). The expected flip rate under independence of the two states was also computed as follows:

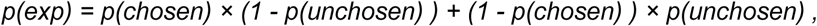

where *p(chosen)* and *p(unchosen)* are the sign-consistency proportions previously derived for each probability state.

### Statistical analyses

Unless noted otherwise, statistical comparisons used linear or generalized linear mixed-effects models (*fitlme* or *fitglme* in MATLAB) with monkey and session as random intercepts. Non-parametric tests (two-sided binomial tests, z-tests) were used where the sampling distribution was not Gaussian, as detailed in the respective sections. Multiple comparisons were corrected using the Benjamini-Hochberg FDR procedure (q=0.05) applied within each analysis family, as described in each section. All analyses were implemented in MATLAB (R2024a/b).

## Data and code availability

The dataset was deposited on Zenodo (https://doi.org/10.5281/zenodo.17524410) and was previously described in London et al (London et al., 2026). Analysis code will be deposited on our lab GitHub (https://github.com/RudebeckLab) before publication.

## Generative AI statement

The authors declare that generative AI was used in the creation of this manuscript. Claude (Anthropic) and ChatGPT (OpenAI) were employed to assist in debugging Matlab code and improving the manuscript readability. All AI-generated content was reviewed and verified by the authors and we take full responsibility for the content of this manuscript.

## AUTHORS CONTRIBUTIONS

Conceptualization and Methodology: FMS, PHR; Investigation, Data Curation: FMS; Analysis, Software and Visualization: FMS, NV; Writing – Original Draft: FMS, NV, PHR; Review/Editing: FMS, NV, PHR; Funding Acquisition and Supervision: FMS, PHR.

## ACKNOWLEDGEMENTS

This work was supported by a National Institute of Mental Health BRAINS award to PHR (R01s MH110822; MH132064), a young investigator grant from the Brain and Behavior Foundation (NARSAD) to PHR, a Philippe Foundation award to FMS, seed funds from the Icahn School of Medicine at Mount Sinai to PHR. We thank members of the Rudebeck lab for comments on the analyses.

